# Dark selection for JAK/STAT-inhibitor resistance in chronic myelomonocytic leukemia

**DOI:** 10.1101/211151

**Authors:** Artem Kaznatcheev, David Robert Grimes, Robert Vander Velde, Vincent Cannataro, Etienne Baratchart, Andrew Dhawan, Lin Liu, Daria Myroshnychenko, Jake P. Taylor-King, Nara Yoon, Eric Padron, Andriy Marusyk, David Basanta

## Abstract

Acquired therapy resistance to cancer treatment is a common and serious clinical problem. The classic U-shape model for the emergence of resistance supposes that: (1) treatment changes the selective pressure on the treatment-naive tumour; (2) this shifting pressure creates a proliferative or survival difference between sensitive cancer cells and either an existing or *de novo* mutant; (3) the resistant cells then out-compete the sensitive cells and – if further interventions (like drug holidays or new drugs or dosage changes) are not pursued – take over the tumour: returning it to a state dangerous to the patient. The emergence of ruxolitinib resistance in chronic myelomonocytic leukemia (CMML) seems to challenge the classic model: we see the global properties of resistance, but not the drastic change in clonal architecture expected with the selection bottleneck. To study this, we explore three population-level models as alternatives to the classic model of resistance. These three effective models are designed in such a way that they are distinguishable based on limited experimental data on the time-progression of resistance in CMML. We also propose a candidate reductive implementation of the proximal cause of resistance to ground these effective theories. With these reductive implementations in mind, we also explore the impact of oxygen diffusion and spatial structure more generally on the dynamics of CMML in the bone marrow concluding that, even small fluctuations in oxygen availability can seriously impact the efficacy of ruxolitinib. Finally, we look at the ability of spatially distributed cytokine signaling feedback loops to produce a relapse in symptoms similar to what we observe in the clinic.

## 1 Introduction

Chronic myelomonocytic leukemia (CMML) is type of leukemia that usually occurs in the elderly and is the most frequent myeloproliferative neoplasm [1]. It has a median survival of 30 months, with death resulting from progression to AML in 1/3rd of cases and cytopenias in the others. In 2011, the dual JAK1/JAK2 inhibitor, ruxolitinib was approved for treatment of the related cancer of myelofibrosis based on its ability to relieve the symptoms of the disease. Recently, it has also started to see use for CMML [2]. Unfortunately, as with most targeted therapies, resistance eventually develops.

The classic model for the emergence of resistance rests on the following microdynamical assumptions: (1) treatment changes the selective pressure on the treatment-naive tumour; (2) this shifting pressure creates a proliferative or survival difference between sensitive cancer cells and either an existing or *de novo* mutant; (3) the resistant cells then out-compete the sensitive cells and – if further interventions (like drug holidays [3] or new drugs [4] or dosage changes [5]) are not pursued – take over the tumour: returning it to a state dangerous to the patient. Clinically this typical process of response and relapse is characterized by a (usually rapid) decrease in tumour burden followed by a transient period of low tumour burden, and finally a quick return of the disease. This is the classic U-shape model of resistance.

When treating CMML with ruxolitinib, clinicians typically see the drastic reduction and then relapse in symptoms (most notably fatigue and spleen size) but none of the microdynamical signs of the classic model of resistance. Merlevede *et al.* [6] used whole-genome sequencing to show that the mutation allele burden and clonal architecture of the bone marrow in CMML remained unchanged during treatment. This suggests that both the total tumour burden and proportional genetic composition in the bone marrow remained static while resistance arose. We see the global properties of resistance, but not the evidence of selection.

This also suggests that it is unlikely that the genotype is acting as the unit of selection for ruxolitinib resistance in myeloid neoplasms. Instead, we have to turn to other units of selection. Units that can propagate epigenetically against an unchanging genetic background. The most obvious candidates are in the cytokine network, which is known to be a factor in the symptoms of CMML. In particular, heterodimeric activation of the JAK-STAT pathway – when the second JAK2 at the cytokine receptor is replaced by a JAK1 or TYK2 [7] – seems like a promising candidate for the unit of selection.

Given the absence of the usual signs of a selective bottleneck, the evolutionary dynamics underlying resistance in CMML are hard to observe. In analogy to physics and recent work on apparently ‘missing’ genes [8], we call this apparently ‘missing’ selective process: dark selection. It is our goal in this report to sketch the question of dark selection for CMML resistance. We achieve this in part by providing some candidate mechanisms for dark selection.

This paper is structured as follows. First, we consider three different alternative effective (cell population level) theories for explaining dark selection: Darwinian (Section 2.1), and two Lamarckian models (Section 2.2): one cell-autonomous and the other non-cell autonomous. These Lamarckian models require a ratcheted phenotypic switch for the proximal resistance mechanisms (like the TYK-bypass described by Koppikar et al. [7]), so we propose a candidate molecular model of this in Section 3.1 as a reductive implementation. Once models shift from an effective description to a reductive implementation, concerns like the spatial structure of the bone marrow become increasingly more important. In Section 3.2, we explore how oxygen is spatially distributed in the bone marrow and how small changes in oxygen penetration and consumption rates can have a large impact on the cells therapy response. This suggest an alternative mechanisms of resistance and/or an amplifier for the others. Finally, in Section 3.3 we consider some of the cytokine signaling network ideas from Section 2.2 in a spatially structured setting. Section 4 summarized the results and sketches future directions.

## 2 Darwinian and Lamarckian population-level models

### 2.1 Hidden Darwinian selection

Drastic population changes are not necessary for evolution. Theoretical and mathematical biologist using evolutionary game theory have traditionally Evolutionary game theorists are more than comfortable thinking about evolution in constant sized populations, and Darwinian evolution itself emerged in the context of a species competing at carrying capacity. As such, the proper null model for us is selection at carrying capacity: a hidden Darwinian selection.

For the hidden Darwinian selection model, we introduced selection through a highly reduced cell turnover. The hypothesis is that as the tumour fills up the bone marrow, it pushes the extra daughter cells out into the peripheral blood; these accumulate in the spleen and cause its drastic enlargement. As therapy takes effect, the division rate of sensitive cells is greatly reduced. The reduction is enough so that many fewer cells are pushed into the periphery, but not so drastic that the tumour burden decreases. Fewer excess cells are made, but the made cells are still excess.

With this model of hidden selection, it is possible to recapitulate the dynamics of the spleen shrinking and relapsing. As we do in figure 1. However, there are tensions with our microdynamical knowledge in the bone. Merlevede *et al.* [6] observed no changes in the clonal architecture of the tumour, meaning that this hidden selection would have to be epigenetic. They also don't see large changes in proliferation rate, which would be required for this model. In future experiments, it would be useful to measure the variance in proliferation rates carefully to rule out hidden selection.

**Figure 1:**
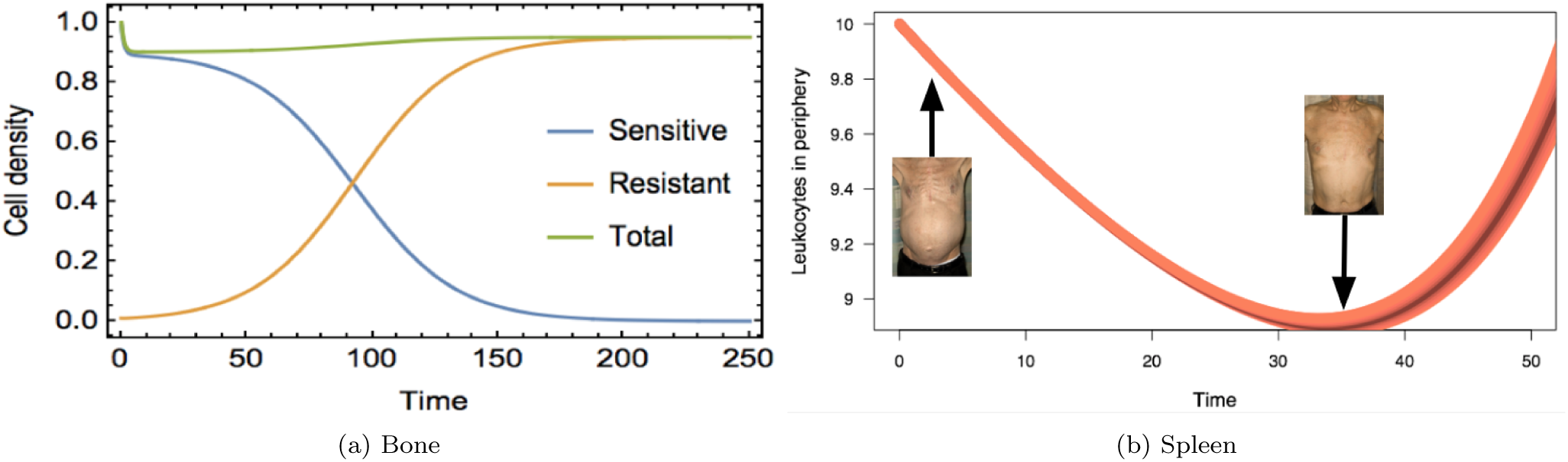
Model of hidden Darwinian selection. (a) A plot showing how Darwinian selection could be acting even when no discernible change in the population size is observed. (b) Even when the tumour size has not changed, the impact of the treatment on the spleen is clearly visible and follows the traditional U-shape.

### 2.2 Lamarckian selection: cell-autonomous and non-cell autonomous

Since we are forced to move from the genetic to the epigenetic level of description, it becomes important to suggest a plausible mechanism for heritable epigenetic effects. One hypothesis is that the acquired JAK-TYK bypass described by Koppikar et al. [7] is heritable. There are many mechanisms by which this non-genetic heritability might be achieved [9] and thus leading to Lamarckian selection. For example, the non-genetic heritability mechanism might be the up-regulation of TYK2 following successful discovery and then Poisson variation in the inheritance of the protein among the two daughter cells. Receptors can be divided in a similar way, probably even more uniformly due to the even division of the cytoplasm. And since daughter cells occupy area close to where their mother cell used to be, local extracellular cytokine concentration are also inherited. We discuss the microdynamical details of this in section 3.1.

At the population level, this allows us to think of the discovery of the heterodimerization bypass as a therapy-induced mutation. The central question becomes if this mutation rate is constant or dependent on the local concentration of cytokines. If it is constant then we have a standard Lamarckian model and if it increases with cytokine concentration then our process is non-cell autonomous. This is represented in Figure 2a, with the Lamarckian process in the red panel and non-cell autonomous in the blue. On the left of each panel is the CMML cell in drug, with the standard pathway blocked. Non-cell autonomous process have been studied in cancer both theoretically – with tools like evolutionary game theory [10-14] – and experimentally [15-17], and are believed to be important in the emergence of resistance in some systems [13, 14, 17]. For the Lamarckian process, their rate of discovery of the heterodimerization bypass is independent of the number of cytokines around them, for the non-cell autonomous it is low with few cytokines and high with many.

**Figure 2:**
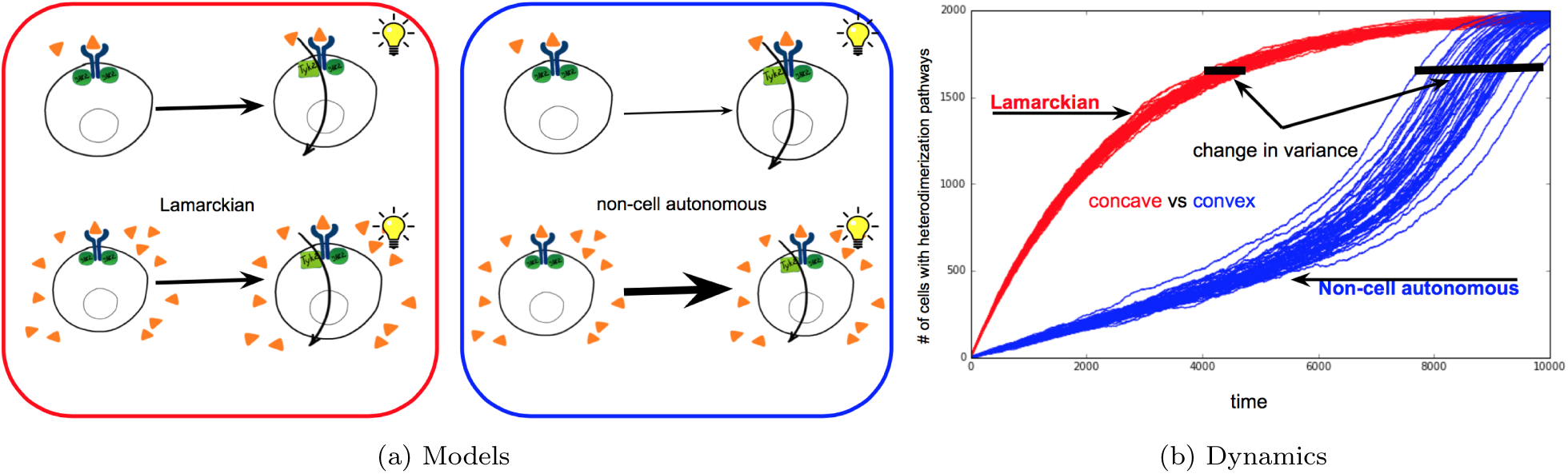
Dynamic models of Lamarckian cell-autonomous (red) vs. non-cell autonomous (blue) dark selection. (a) Discovery of JAK-TYK bypass is independent of levels of cytokines in the cell-autonomus (red) case or creates a positive-feedback loop in the non-cell autonomous (blue) case: as more cells discover the bypass to more cytokines, other cells are more likely to discover the same bypass. (b) This results in qualitative different dynamics: cell-autonomous process has a concave path with relatively low variance compared to non-cell-autonomous process with a convex (or sigmoidal) path with higher variance.

For the dependence of the JAK-TYK bypass discovery rate on the levels of cytokines in the non-cell autonomous model, it is again good to turn to models at the levels of receptors. Depending on the sort of feedback loops possible within the cell's internal signaling, it is possible to get different functional forms for the mutation rate. To avoid testing against any possible mutation function, it is best to consider the functional form that comes out of biologically reasonable assumptions of the cellular pathways. We propose a specific biological basis in Section 3.1. Depending on the sort of data available, it might be worth considering adapting recent techniques like Lever *et al.* [18] for inferring the form of the minimal signaling pathway from standard molecular and systems biology experiments.

As seen from Figure 2b, these two processes result in drastically different relapse curves. In red is the Lamarckian process, and in blue is the non-cell autonomous. Both have the same average mutation rate, but for the former it is constant through time while the latter scales with the amount of cytokines and thus increases over time. The result is that the relapse curve for blue is much more convex and has a higher variance.

With qualitative different curves like the ones above, we can hope to distinguish between the models with the sort of noisy data that one can expect from biological experiments. In particular, from proximity ligation of JAK-TYK, one could see which cells have discovered the bypass in bone histologies of mice. By taking bone histologies from an experimental model like mice sacrificed at different time points, we can build up a time series of acquisition of resistance to test our models against.

## 3 Ratcheting & microenvironment

### 3.1 Ratcheting the TYK-bypass

Whereas the above is an effective theory at the level cell populations – akin to a game assay [17] – for distinguishing between cell-autonomous and non-cell autonomous processes, it (a) Homeostasis (b) Switch discovery can also be useful to have a reductive theory at the level of molecules and pathways. We need to find a stochastic ratcheted phenotypic switch among the pathways of the CMML cells. Here, the Koppikar *et al.* [7] study of the microdynamical basis of roxlitinib resistance becomes very valuable. In the untreated cancer cell, two JAK2 attach to a receptor, cross-phosphorylate each other, and then active a STAT. Roxlitinib blocks the ability of the JAK2 pair to do this, shutting down the JAK-STAT pathway. Koppikar et al. [7] showed that some CMML cells find a way around this by having JAK2 heterodimerize with JAK1 or TYK2 instead of another copy of JAK2. Once the cell finds this alternative pathway, it is able to maintain it and start over-producing cytokines as it did before. With enough cells discovering this bypass pathway, global cytokine levels can become elevated again, leading to a return of symptoms.

To discover a reaction that explains our higher level models it is prudent to look for the simplest possible reaction that fulfills the requirements of these models, then try to break down actual biological reactions in CMML towards our reductionist model to see if they can fulfill these requirements. As such we first consider one of the simplest reactions possible, where one type of chemical transforms into another: A ↔ *B*. One can work out that a stable equilibria occurs at 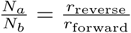 (where *N*_a_ is the number of A molecules, *N_b_* is the number of *B* molecules, *r_forward_* is the reaction rate as A transforms to *B* and *r_reverse_* is the reaction rate as *B* transforms to A).

Our model requires a chemical reaction that can switch from one state to another either randomly or due to the migration of molecules from a nearby cell. With an unstable equilibrium we create a barrier which can be crossed in either of the previous two scenarios, but ordinarily would not. We therefore split our simpler reaction into several parts to create this unstable equilibrium. We start with a heterodimerization reaction where A and B bind: *A* + *B* → *AB*.

If we then converted AB to two *B*’s we would have a classic autocatalysis reaction. But that is not enough for our purposes since this reaction relies just as much on A as it does on *B*: we need the reaction to rely more on B since unstable equilibria rely on the predominance of the species being created; this is also why we need autocatalysis.

Therefore, we need a “peer-pressure” reaction step: AB + *B* → ABB → 3B. Here A is outnumbered by *B* in the complex and is therefore converted to *aB.* We do the same for *B*’s converting to *A’s*: 3A ← *AAB* ← *AB* + *A*.

Note that the net reaction is still one A being converted to one *B*. If we assign reaction coefficients of 1 to all reactions except those where complexes are used up (which will have coefficients that are >> 1 (here 1000) in order to keep the concentration of complexes small) we can approximate the change in the expected number of *B* molecules as:

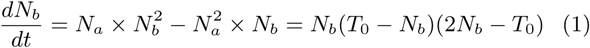

where *T*_o_ = *N*_*a*,0_ + *N_b,o,_* the total number of *A* and *B* at the beginning of the simulation.

For this sytem, three equilibria exist: one with all A, one with all *B* and one in between where 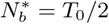; by checking the signs on both sides of this last equilibrium we see that it is unstable.

If we plug these values into the Gillespie algorithm the probability of having an A turn into a *B* when there are no other *B’s* around is 0. Therefore, in order to have a stochastic change in state, we need a leak equation where the net reaction (*A* ↔ *B*) occurs spontaneously, but at a lower rate than the autocatalytic reaction.

These reactions act as both ‘mutation’ and mode of inheritance. Stochastic changes in the amount of chemical due to the ‘leak’ reactions can lead to a spontaneous change in state in some cells – mutation (as seen in figure 3b but not in figure 3a). But other cells can also impose this state by increasing the number of molecules of one chemical species, either through diffusion or through activation by some other diffusing species: inheritance. Hence two requirements, variation and inheritance, can be satisfied by a relatively simple set of chemical reactions.

**Figure 3:**
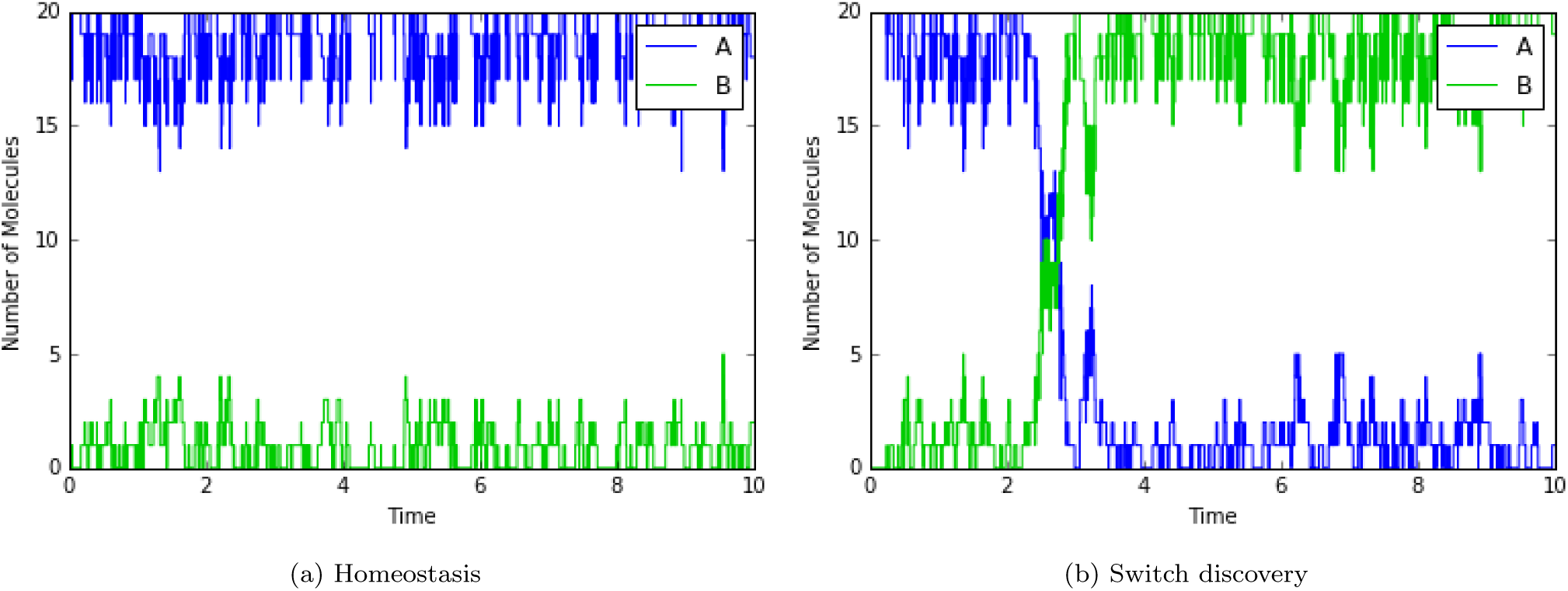
Dynamics of a chemical reaction network with possibility of ratcheted switch [19]. Due to stochastic fluctuations, even with the same initial conditions, it is possible to either (a) not discover the JAK-TYK switch or (b) to ratchet into the alternative pathway (b).

### 3.2 Role of oxygen in the bone marrow

Oxygen distributions inside tumour micro-environments are typically complex, with pockets of chronic hypoxia being a common facet in solid tumours [20–23]. The physical rationale for this is relatively straightforward - as oxygen diffuses through tissue, it is depleted by cells as a vital component in aerobic respiration [24–26]. In healthy tissue, there is generally adequate vascular supply of oxygen and no hypoxia. However, the tumour micro-environment is much more chaotic, marked by tortuous and poorly perfused vascular networks with an excess of cells.

As a consequence, oxygen supply is generally disordered and demand higher, leading to regions of extensive hypoxia. Whilst the mechanism might be simple, the net result is intrinsically complex. This diffusion-limited hypoxia gives rise to a number of detrimental effects. It has also long been known that extensive hypoxia has a negative implication for patient prognosis [27]. The reason for this is deeply evolutionary; under hypoxic conditions, tumour cells can respond to hypoxia by activating oxygen sensitive signalling pathways [28], including hypoxia inducible factor pathways and the unfolded protein response [29].

While the precise mechanisms remain poorly understood, it is thought that these signaling pathways alter gene expression in an attempt to promote survival under adverse conditions, and ultimately allow cellular phenotypes to arise with evasive mutations, including the ability to metastasize. This is of importance for solid tumours, where spatial limitations on oxygen diffusion yield such chaotic pockets of anoxia.

It might even be tempting to dismiss it as a dead-end avenue. But given the ubiquity of micro-environmental influence, we opted to investigate it further in the search for potential mechanisms of ruxolitinib resistance, or at the very least to understand what factors might influence this. So if we were to search for the mechanistic fingerprint of oxygen effects, where better to look than the scene of the crime – inside the bone marrow itself.

Bone marrow is relatively unusual in one respect, with a surprisingly low oxygen partial pressure, ranging typically from a low of 9 mmHg to highs of 32 mmHg [30]. To see this in perspective, typical partial pressures at the capillary wall tend to be around 40 mmHg, whereas many human organs have oxygen tensions well in excess of 150 mmHg. It is important to note that despite this low relative oxygen tension, bone narrow is not hypoxic in the conventional sense. Cells can quite happily survive much lower oxygen pressures, happily undergoing mitosis as low as 0.5 mmHg [31, 32].

Let’s look again at how ruxolitinib itself works. The drug doesn’t attack the cancer – ruxolitinib is a Janus kinase inhibitor, and works by repressing the JAK-STAT pathway to ease the symptoms of the illness.

If the oxygen micro-environment interfered with the drug rather than the CMML cells, then perhaps these seemingly disparate facts might tell us another story. As ruxolitinib’s mechanism of action relies on its ability to block JAK-STAT signally, we might ask whether availability of oxygen potentially modulates or diminishes inhibitor efficacy. Wang *et al.* [33] had looked at the levels of JAK-STAT signalling under both low and high oxygen conditions. And in both cases, JAK-STAT signalling was increased under hypoxia. It’s important to note that the paper didn’t use ruxolitinib specifically but instead a different JAK-STAT inhibitor, AG490, but for our purposes we initially assume the mechanisms are broadly similar. Hypoxia diminished the effectiveness of the inhibitor drastically, and this reduction impact was non-linear and markedly less effective when oxygen was scarce.

This finding was intriguing, suggesting that oxygen might indeed have a role, albeit an unusual one. If treatment was to fail, there mightn’t be any need for clinical hypoxia – just a small reduction in available oxygen tension might be enough to hobble the drug’s ability to quench JAK-STAT signals. But for this to be the case there would have to be something capable of changing the oxygen profile in the bone marrow. This hypothesis could have at least two mechanistic explanations– the first is simply that increased density of cells depletes available oxygen more than in non-cancerous marrow. Another potential reason is the strange metabolistic properties of tumour cells – oxygen consumption rate (OCR) has a distinct effect of available oxygen, and if cells had up-regulated their OCR this could easily give rise to diminished oxygen availability.

Both these mechanisms are plausible. But if this were the potential mechanism of treatment resistance, then the reduction in oxygen would have to be relatively subtle – enough that to impact the drug efficacy without significantly stressing the cells to the point where they would display hallmarks of clinical hypoxia. To look into this, we used a vascular model based on Grimes et al. [32] to simulate the oxygen profile between two hypothetical blood vessels, each with a wall-pressure of 32 mmHg. We then changed the OCR of the bulk tissue slightly, giving rise to subtly different oxygen profiles (mean p02 15.9 mmHg, 13.1 mmHg and 10.1 mmHg respectively), shown in figure 4a. Crucially, in all simulated cases the global oxygen pressure is far beyond that which would be expected to have clinical significance.

**Figure 4:**
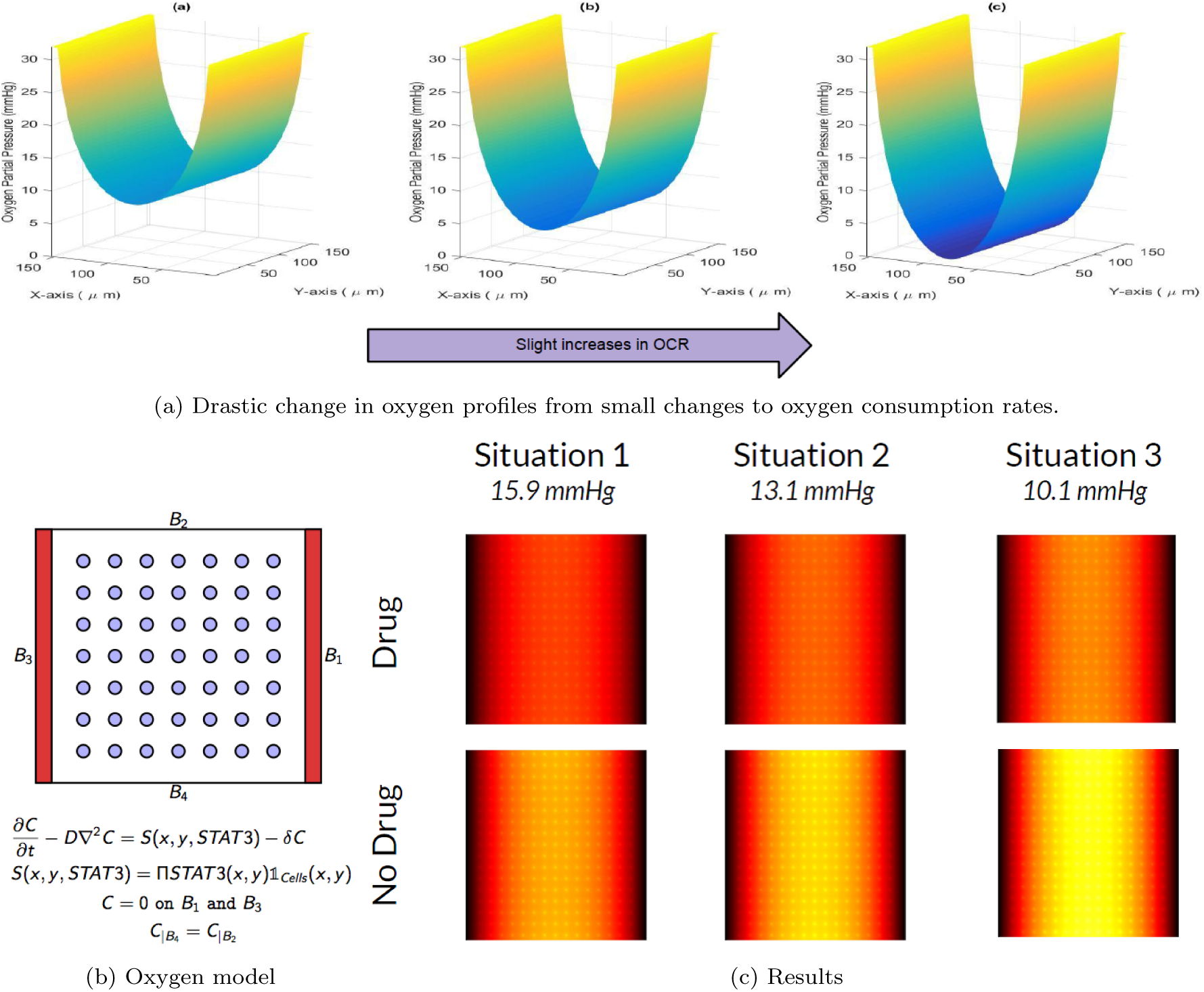
Modeling oxygen and the effectiveness of drug on the JAK-STAT pathway. (a) Drastic decrease in oxygen partial pressure in the bone marrow from mild increase in oxygen consumption rate. (b) PDE model of STAT activity and its interaction with the oxugen and cytokine gradient. (c) Resulting levels of cytokine activity in the bone marrow with changes in the oxygen partial pressure. Bright corresponding to high activity.

Armed with a simulated oxygen profile and the experimental data on the variation of JAK-STAT inhibition with oxygen tension, we produced a simplified 2D model of how these effects might interplay. In the bone marrow, ruxolitinib diffused in from the blood vessels, across a discrete matrix of cells with varying oxygen tension, each of which produced JAK-STAT signal. In the model, We assumed that the drug perfectly perfused the tissue and thus the drug level is assumed to be spatially homogeneous. The drug acted to reduce the JAK-STAT signal in a manner proportional to oxygen tension, with the linear coefficient calibrated from experimental data ??. A simplified schematic is shown in figure 4b, alongside the general form of the model PDE. The PDE describes how the concentration of cytokines produces by CMML cells is distributed (by stationary 2D diffusion) over the bone marrow.

With this schema in place, we were free to run a simulation – and the results were striking. Figure 4c indicates that even marginal differences in oxygen profile and mean pO2 had substantial impact of the efficacy of the drug. Indeed, the situation where the drug was applied at a mean pO2 of 10.1 mmHg was virtually identical to an absence of drug at a slightly higher oxygen tension (mean pO2 = 15.9 mmHg). This was a fascinating result, and consistent with the candidate hypothesis. If this were the case, then slight variability in oxygen tension could have a marked effect on treatment, and might be part of the reason for the emergence of resistance.

Of course, this remains at the moment just an interesting candidate hypothesis — we would need a lot more evidence to state that this is the mechanism responsible. What’s difficult to account for with this approach is the timing of and variance in relapse. But it is also an approach that could be combined with any of the previous suggestions as a way to amplify the effect of small epigenetic or signaling changes.

### 3.3 Emergent spatially-explicit signaling network behavior

Inhibition of JAK-STAT signaling axis in leukemic and stromal cells impacts abundance of cytokines, including direct targets of STAT transcriptional regulators, as well as those induced as the result of altered cell-cell communication. The net impact is lower level of activating signaling stimulation in CMMI cells. However, cell signaling is a complex dynamic process involving positive and negative regulatory loops [34]. We speculated that relapse of activation of CMML cells despite the lasting primary effects of JAK inhibition might reflect adaptation of signaling networks operating between cells, perhaps reflecting homeostatic buffering dynamic mechanisms.

To test for the potential feasibility of this hypothesis, we constructed an agent-based model of cell cytokine release to investigate the plausibility that epigenetic and/or phenotype switching to a drug insensitive state may lead to recurrent tumor dynamics. We also use this model to investigate potential differences in dynamics that may arise depending on whether the rate of insensitivity switching is conditional on the local cytokine load. Briefly, our model is initiated with a specified density of immobile cells arranged *in silico* on a 2-dimensional grid. Cells may be active, and release cytokines at every iteration of the simulation, and become inactive if cytokines build up over a specified inactivation threshold. Inactivated cells may become activated after a cool-down period if their local cytokine levels build up over a specified activation threshold. Cytokines diffuse and decay in this space. A drug is introduced uniformly throughout the grid at some time point, and this drug blocks all cytokine release and cytokine response. Due to the persistent lack of cytokines, all cells in the simulation build up a lower tolerance to any cytokines present, i.e., cells have a lower threshold for activation if they come into contact with cytokines, perhaps through known mechanisms such as increased cytokine receptor densities per cell. Cells have some probability to switch to a less sensitive state and release and respond to cytokines even with the drug present. These dynamics reveal that cytokine levels, and therefore tumor activity, return to before-drug levels over time, even with no change in tumor burden (Figure 5).

**Figure 5:**
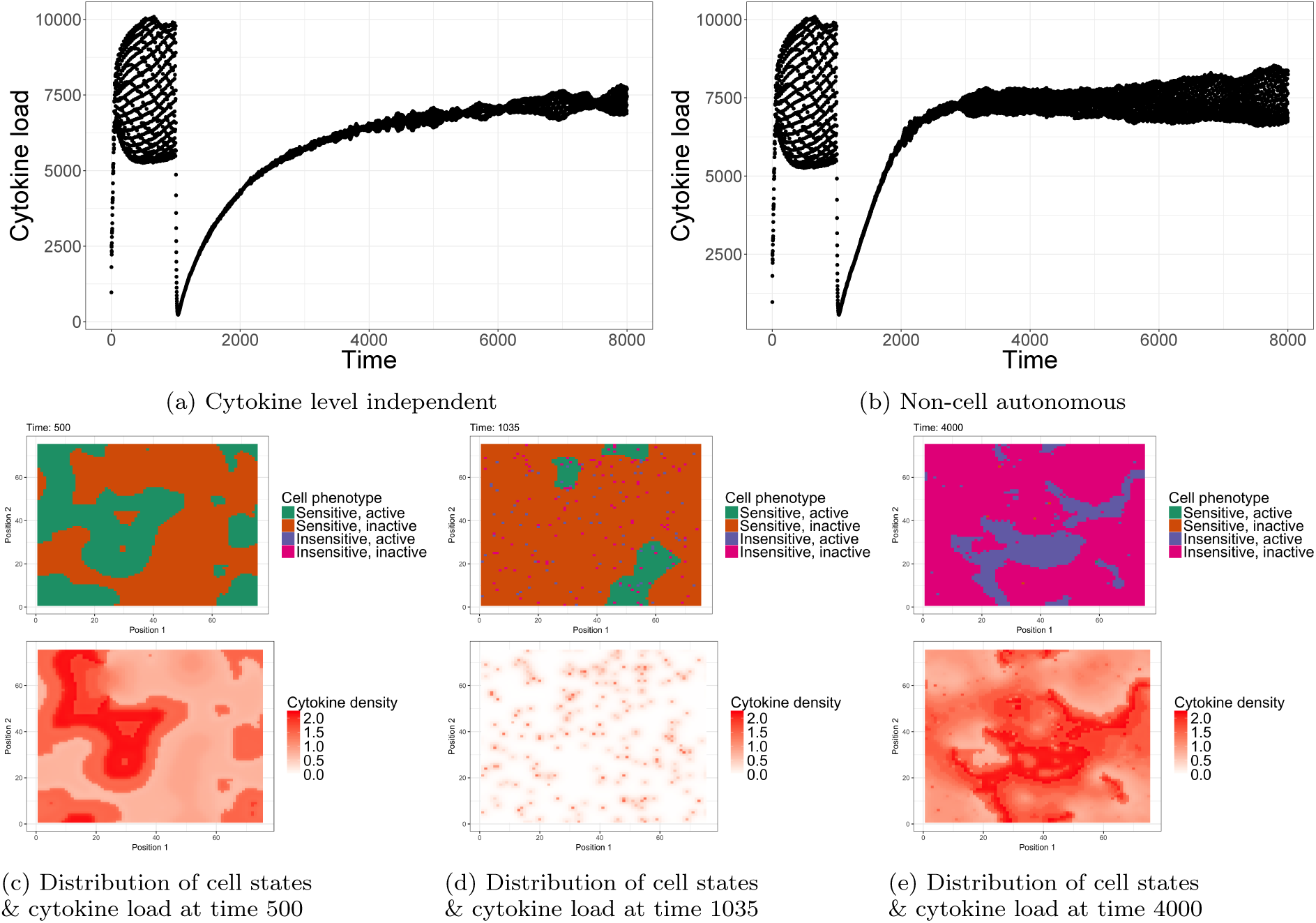
Spatially-explicit signaling network dynamics. After drug is added at time 1000, cells may randomly switch to a drug-insensitive phenotype, and become active and release cytokines. We model scenarios where the probability that a cell becomes drug-insensitive is (a) independent and (b) dependent on local cytokine levels. Local spatial dynamics of cytokine load and cell phenotype states for the cytokine-dependent scenario are shown for time-points (c) before drug, (d) immediately after drug introduction, and (e) after drug has been present for a sufficient amount of time for cytokine level relapse.

We find that cytokine levels return to an equilibrium similar to that found before treatment in both cytokine-level-independent (Figure 5a) and cytokine-level-dependent (non-cell autonomous; Figure 5b) insensitivity scenarios, however, when the rate at which cells switch to drug insensitivity is dependent and positively correlated with the local cytokine loads, the return to equilibrium is faster. For both scenarios, cytokine levels begin low, as the simulations begin with zero cytokines present and a random subset of cells active, and gradually build up to equilibrium (e.g., Figure 5c for the non-cell autonomous scenario). After drug introduction, the cumulative level of cytokines decay to near-zero, as they are no longer produced except for in a small number of cells that entered into a drug-insensitive state (Figure 5d). As time progresses, more cells stochastically become drug-insensitive, and the cytokine levels increase to an equilibrium near that of the pre-drug state of the system (Figure 5e).

## 4 Conclusion

It is hard to solve a problem that almost no one sees. As such, our goal in in this paper was to illuminate an interesting theoretical problem: dark selection. Dark selection is the emergence of resistance without the apparent microdynamical hallmarks of the classic U-shaped model of resistance. We used CMML as our case study. However, we expect that once we learn how to look for it, dark selection might reveal itself in other cancers.

To more clearly define the boundaries of this theoretical problem we proposed some models as potential solutions. We considered two types of models: microdynamical and effective. The goal of the microdynamical models is to propose concrete biological mechanisms at the molecular and micro-environmental level like ratcheting via JAK-TYK bypass, oxygen diffusion affecting the JAK-STAT pathway, and spatial structure of cytokine signaling. These models are meant as heuristics to motivate further discussion with experimentalists. The effective models, however, aimed to connect directly to the type of data that can be gathered from patients, mice, or in-vitro experiments. As such, they sacrifice mechanistic detail to differentiate between three different kinds of selection: hidden Darwinian selection, and Lamarckian selection that is either cell-autonomous or non-cell autonomous. Although these population-level models might not provide insights to the molecular basis for drug targets, they do provide an explicit target population-dynamic for more reductive models to produce.

At the current stage, the goal of these models is not rigour but the encouragement of creative exploration of hypotheses. Our aim is not to solve the problem that we saw, but to use these avenues of answers as a better outline of the terrain. The next step is to design experiments that can help inform these models. By understanding dark selection from both a mathematical and experimental perspective, we will have a new tool in our arsenal against the emergence of therapy resistance in chronic myelomonocytic leukemia and other cancers.

